# Characterization of cellular, biochemical and genomic features of the diazotrophic plant growth-promoting bacterium *Azospirillum* sp. UENF-412522, a novel member of the *Azospirillum* genus

**DOI:** 10.1101/2021.05.06.442973

**Authors:** Gustavo L. Rodrigues, Filipe P. Matteoli, Rajesh K. Gazara, Pollyanna S. L. Rodrigues, Samuel T. dos Santos, Alice F. Alves, Francisnei Pedrosa-Silva, Isabella Oliveira-Pinheiro, Daniella Canedo-Alvarenga, Fabio L. Olivares, Thiago M. Venancio

**Affiliations:** Laboratório de Química e Função de Proteínas e Peptídeos, Centro de Biociências e Biotecnologia, Universidade Estadual do Norte Fluminense Darcy Ribeiro (UENF), Brazil; Núcleo de Desenvolvimento de Insumos Biológicos para a Agricultura (NUDIBA), UENF, Brazil; Laboratório de Biologia Celular e Tecidual, Centro de Biociências e Biotecnologia, UENF, Brazil

**Author notes:** **Corresponding authors:** Thiago M. Venancio; Laboratório de Química e Função de Proteínas e Peptídeos, Centro de Biociências e Biotecnologia, UENF; Av. Alberto Lamego 2000, P5 / sala 217; Campos dos Goytacazes, Rio de Janeiro, Brazil. Fabio L. Olivares: Laboratório de Biologia Celular e Tecidual, Centro de Biociências e Biotecnologia, UENF, Brazil. Contributed equally to this work (Gustavo L. Rodrigues, Filipe P. Matteoli).

## Abstract

Given their remarkable beneficial effects on plant growth, several *Azospirillum* isolates currently integrate the formulations of various commercial inoculants. Our research group isolated a new strain, *Azospirillum* sp. UENF-412522, from passion fruit rhizoplane. This isolate uses carbon sources that are partially distinct from closely-related *Azospirillum* isolates. Scanning electron microscopy analysis and population counts demonstrate the ability of *Azospirillum* sp. UENF-412522 to colonize the surface of passion fruit roots. *In vitro* assays demonstrate the ability of *Azospirillum* sp. UENF-412522 to fix atmospheric nitrogen, to solubilize phosphate and to produce indole-acetic acid. Passion fruit plantlets inoculated with *Azospirillum* sp. UENF-41255 showed increased shoot and root fresh matter, as well as root dry matter, further highlighting its biotechnological potential for agriculture. We sequenced the genome of *Azospirillum* sp. UENF-412522 to investigate the genetic basis of its plant-growth promotion properties. We identified the key *nif* genes for nitrogen fixation, the complete PQQ operon for phosphate solubilization, the *acdS* gene that alleviates ethylene effects on plant growth, and the *napCAB* operon, which produces nitrite under anoxic conditions. We also found several genes conferring resistance to common soil antibiotics, which are critical for *Azospirillum sp.* UENF-412522 survival in the rhizosphere. Finally, we also assessed the *Azospirillum* pangenome and highlighted key genes involved in plant growth promotion. A phylogenetic reconstruction of the genus was also conducted. Our results support *Azospirillum sp.* UENF-412522 as a good candidate for bioinoculant formulations focused on plant growth promotion in sustainable systems.

## INTRODUCTION

Modern agriculture production strongly relies on synthetic fertilizers and pesticides to achieve high productivity levels, which are often a matter of environmental and health concerns [1]. The development of bioinoculants has been considered an alternative to reduce the use of such synthetic compounds [2]. These inoculants typically have one or more strains of plant growth-promoting rhizobacteria (PGPR), which compose a heterogeneous group of bacteria that exert mutualistic interactions with plants [3], often playing key roles in the root microbiome [4]. The rhizosphere is the fraction of soil directly influenced by roots exudates [5], which modulate bacterial diversity across the endorhizosphere, rhizoplane and ectorhizhosphere (reviewed by Ahkami et al. [6]). Several *Azospirillum* isolates have been reported as PGPR, typically as soil-borne bacteria that are competent rhizosphere colonizers [7, 8].

The genus *Azospirillum* belongs to the *Rhodospirillaceae* family, which is mainly constituted of aquatic genera. Nevertheless, Azospirilla are mostly soil bacteria that coevolved with vascular plants [9]. Following its original description [10], multiple plant-associated *Azospirillum* species have been shown to perform direct nitrogen fixation [11, 12], phosphate solubilization [13], drought and salt stress alleviation [14, 15], root development promotion [16], among other processes [17]. These desirable properties have led several *Azospirillum* strains to be used as part of commercial soil inoculants [18]. Genes associated with these features have been identified in several publicly available *Azospirillum* genomes, such as *nif* [19], ACC-deaminase (*acds*) [20, 21], *pqq* [22], and indole acetic acid biosynthesis genes (e.g., *iaaH*, *iaaM* and *ipdC*) [23, 24].

Recently, our research group has isolated several plant growth-promoting bacterial strains from vermicompost [25, 26] and from the rhizosphere of tropical fruit trees. Among these strains, *Azospirillum sp.* UENF-412522 was isolated from passion fruit rhizoplane. Here we report *in vitro* and *ex vitro* plant growth promotion capabilities, biochemical tests, microscopy analysis of root colonization, whole-genome sequencing, and comparative genomic analyses to investigate the plant growth promotion properties of this isolate, as well as to uncover genes involved in other ecophysiological processes. Together, our results support the potential of *Azospirillum sp.* UENF-412522 to be used in bioinoculant formulations.

## RESULTS AND DISCUSSION

### Identification of the isolate and assessment of plant growth promotion properties

The strain UENF-412522 was isolated from passion fruit rhizoplane and formed a pellicle typical of diazotrophic bacteria on N-free semisolid NFb medium. It is a Gram negative, slightly curved rod-shaped bacterium with apparently no polymorphic 1.2 × 0.7 μm cells. In order to characterize the isolated strain, a fragment of the 16S rDNA gene was amplified by polymerase chain reaction (PCR); this sequence was deposited in Genbank under the accession KU836626.1. The 16S rDNA maximum likelihood (ML) phylogenetic reconstruction (Figure S1) has not allowed us to classify the strain at the species level. Nevertheless, it is clear that it belongs to the *Azospirillum* genus, leading us to name it *Azospirillum sp.* UENF-412522.

Given the evidence supporting *Azospirillum* sp. UENF-412522 as a novel *Azospirillum* species, we used an API 50 CH system and found that it grows on 18 out of 49 tested carbon sources: glycerol, D-arabinose, L-arabinose, D-ribose, D-xylose, methyl-beta-D-xylopyranoside, D-galactose, D-glucose, D-fructose, D-mannose, L-sorbose, amygdalin, aesculin ferric citrate, D-melibiose, glycogen, xylitol, D-xylose, D-fucose. On the other hand, this strain was unable to grow on: erythritol, L-xylose, D-adonitol, L-rhamnose, dulcitol, inositol, D-mannitol, D-sorbitol, Methyl-alpha-D-mannopyranoside, Methyl-alpha-D-glucopyranoside, N-acetylglucosamine, arbutin, salicin, D-cellobiose, D-maltose, D-lactose (bovine origin), D-saccharose (sucrose), D-trehalose, inulin, D-melezitose, D-raffinose, amidon (starch), gentiobiose, D-turanose, D-tagatose, L-fucose, D-arabitol, L-arabitol, potassium gluconate, potassium 2-ketogluconate, potassium 5-ketogluconate. Comparison of carbon utilization patterns of *Azospirillum* sp. UENF-412522 and those of the type strains of *Azospirillum lipoferum* ATCC 29707 [10], *Azospirillum doebereinerae* DSM 13131 [27] and *Azospirillum brasilense* ATCC 29145 showed that four C-sources (i.e. D-ribose, D-mannitol, D-sorbitol and N-acetylglucosamine) could be used to discriminate these isolates (Table S1).

Next, we performed *in vitro* tests that would account for the ability of *Azospirillum* sp. UENF-412522 to promote plant growth. The N_2_-fixation was supported by positive growth on N-free semisolid medium and subsequently confirmed by the amplification of *nifH* (Figure 1A). We also confirmed the nitrogenase activity by using the acetylene reduction assay, which showed a rate of 28.3 ± 5.1 nmol.h^−1^ ethylene (Figure 1B).

**Figure 1:**
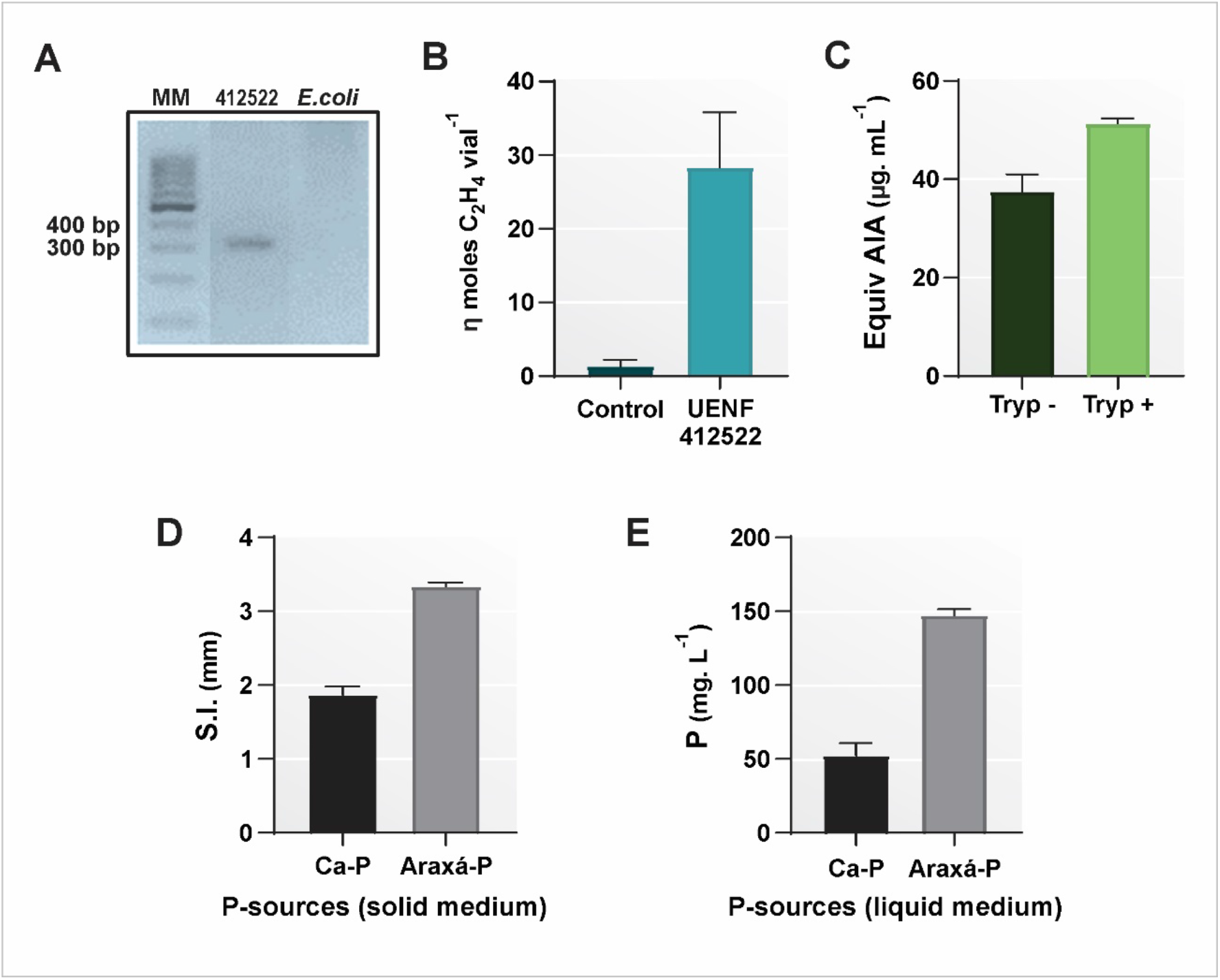
Plant growth promotion *in vitro* traits of *Azospirillum* sp. UENF-412522. (A) Amplification of *nifH* using *E. coli* as negative control; molecular weight marker (MM). (B) Nitrogenase activity by the acetylene reduction assay. (C) Indole acetic acid (IAA) production with and without L-tryptophan. Araxá P-rock and calcium phosphate solubilization in solid (D) and liquid (E) medium.

Indole-acetic acid (IAA) production was investigated in DYGS medium, with and without L- tryptophan, resulting in IAA concentrations of 51.34 ± 1.1 μg.mL^−1^ and 37.35 ± 3.5 μg.mL^−1^, respectively (Figure 1C), supporting the capacity of *Azospirillum* sp. UENF-412522 to synthesize this important plant hormone.

We also tested whether *Azospirillum* sp. UENF-412522 is able to solubilize Araxá P-rock and calcium phosphate in solid medium. We found that this strain formed a halo with a solubilization index (SI) of 3.33 ± 0.05 and 1.87 ± 0.11 for Araxá P-rock and calcium phosphate, respectively (Figure 1D). We confirmed the P solubilization activity using a liquid medium assay, which showed 147.5 ± 3.9 mg.L^−1^ and 52.2 ± 8.6 mg.L^−1^ solubilized P from Araxá P-rock and calcium phosphate (Figure 1E), respectively. Conversely, zinc solubilization was not detected, likely because of zinc sensitivity in this strain.

The plant growth-promoting effects of a given bacterial strain often result from several concerted processes. Therefore, we inoculated passion fruit plantlets with *Azospirillum* sp. UENF-412522 and found a clear growth increment in inoculated versus non-inoculated plantlets. After 10 days, inoculated plantlets had a significant increase in height (54.5%), root fresh matter (88.6%), root dry matter (61.4%), shoot fresh matter (13.8%) and root length (40%). No significant difference was found for shoot dry matter (Figure S2).

Under gnotobiotic conditions, inoculated plantlets showed a larger diazotrophic bacteria population associated with rhizosphere, rhizoplane, and roots (Figure S3). For rhizosphere and rhizoplane compartments, the population density was significantly greater than control plantlets (approximately 1.5 log_10_ bacteria cell per g of soil or root). The bacterial population inside the plantlets showed no difference between inoculated and non-inoculated plantlets, further supporting that *Azospirillum* sp. UENF-412522 preferentially colonizes the rhizosphere/rhizoplane. In parallel, samples from seedlings inoculated with *Azospirillum* sp. UENF-412522 at 1 and 7 days after germination (d.a.g.) were viewed under SEM (Figure 2), confirming the root surface as the preferential colonization site. After radicle emergence, single cells were noted at the bottom region of the radicle (Figure 2A), and bacteria aggregates can be seen in the vicinity of the crack zone between the radicle and the tegument (Figure 2C-D). Interestingly, sparse stomata-like structures were seen at the radicle, which seems to be an infection point for the bacteria (Figure 2B). Bacteria cells interact with the root surface mainly by apolar attachment and by fibrillar material anchoring the plant cell-wall, establishing an aggregate network (Figure 2E). At seven d.a.g., bacteria successfully colonized the root hair zone (Figure 2F) and the elongation/differentiation zone (Figure 2G) of the main developed root.

**Figure 2:**
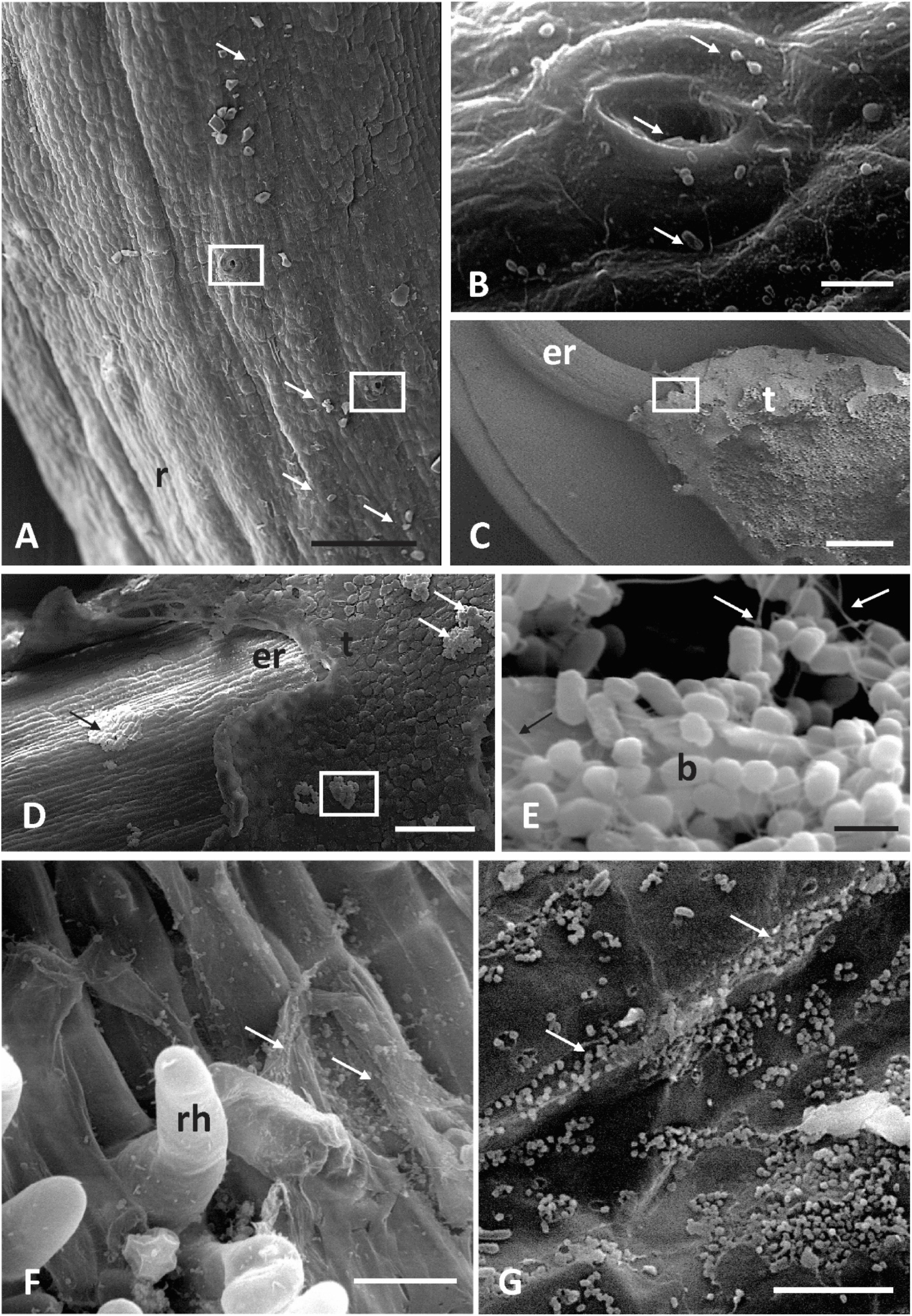
Scanning electron microscopy (SEM) of passion fruit (*Passiflora edulis* subsp. *edulis*) seed to seedlings phase inoculated with *Azospirillum* sp. UENF-412522 in an axenic cultivation system at one day (A-E) and seven days after germination (F-G). (A) General view of the emerged radicle surface (r) with single bacteria cells attached (arrows). Note the sparse rudimentary stomata (rectangle magnified in B), bar = 100 μm; (B) Bacteria cells in the vicinity and into the stomata pore (arrows), bar = 5μm; (C) General view of the emerged radicle (er) breaking the seed tegument (t), rectangle magnified in D, bar = 1000 μm; (D) Transition zone between broken tegument (t) and emerged radicle with bacterial aggregates visible in surface tegument (white arrows) and emerged roots (black arrow), rectangle magnified in E, bar = 100 μm; (E) Bacteria aggregates (b) attached by fibrils in the seed surface (black arrow) and linking cell-to-cell (white arrows), bar = 1 μm; (F) Bacteria cells colonizing the root hair (rh) zone as a single-cell by apolar attachment, bar = 20 μm; (G) Aggregated bacteria cells colonizing the elongation-differentiation root zone mainly by apolar attachment and associated to the epidermal cell-wall-junction, bar = 10 μm.

### Genome assembly and strain identification

The promising *in vitro* and *ex vitro* results prompted us to sequence the *Azospirillum sp.* UENF-412522 genome. We used an Illumina HiSeq 2500 instrument (paired-end mode, 2 × 100 bp reads). Sequencing reads were quality filtered and assembled with SPAdes (see methods for details). The assembly consists of 101 contigs (length ≥ 500 bp), encompassing 7,360,543 bp, with a 67.92% GC content, N50 and L50 of 175,376 bp and 13 respectively. The genome harbors 6,508, 71 and 8 protein-coding, tRNA and rRNA genes, respectively.

Genome relatedness within the *Azospirillum* genus was computed using the average nucleotide identity (ANI) and digital DNA-DNA hybridization (dDDH) of all *Azospirillum* spp. genomes deposited in RefSeq (n = 48). ANI values were used to build a genome-to-genome distance matrix and a neighbor-joining dendrogram (Figure 3A). This analysis clearly shows distinct clusters comprising *A. brasilense* and *A. tiophilum* isolates. In contrast, none of the remaining strains formed a consistent clade. Further, *A. lipoferum* RC and 4B isolates did not cluster together, indicating that they do not belong to the same species. We found greater genetic diversity in the isolates that do not belong to these clusters (Figure 3B). Among those is *Azospirillum* sp. UENF-412522, which does not cluster with any other known strain, suggesting that it represents a novel *Azospirillum* species.

**Figure 3:**
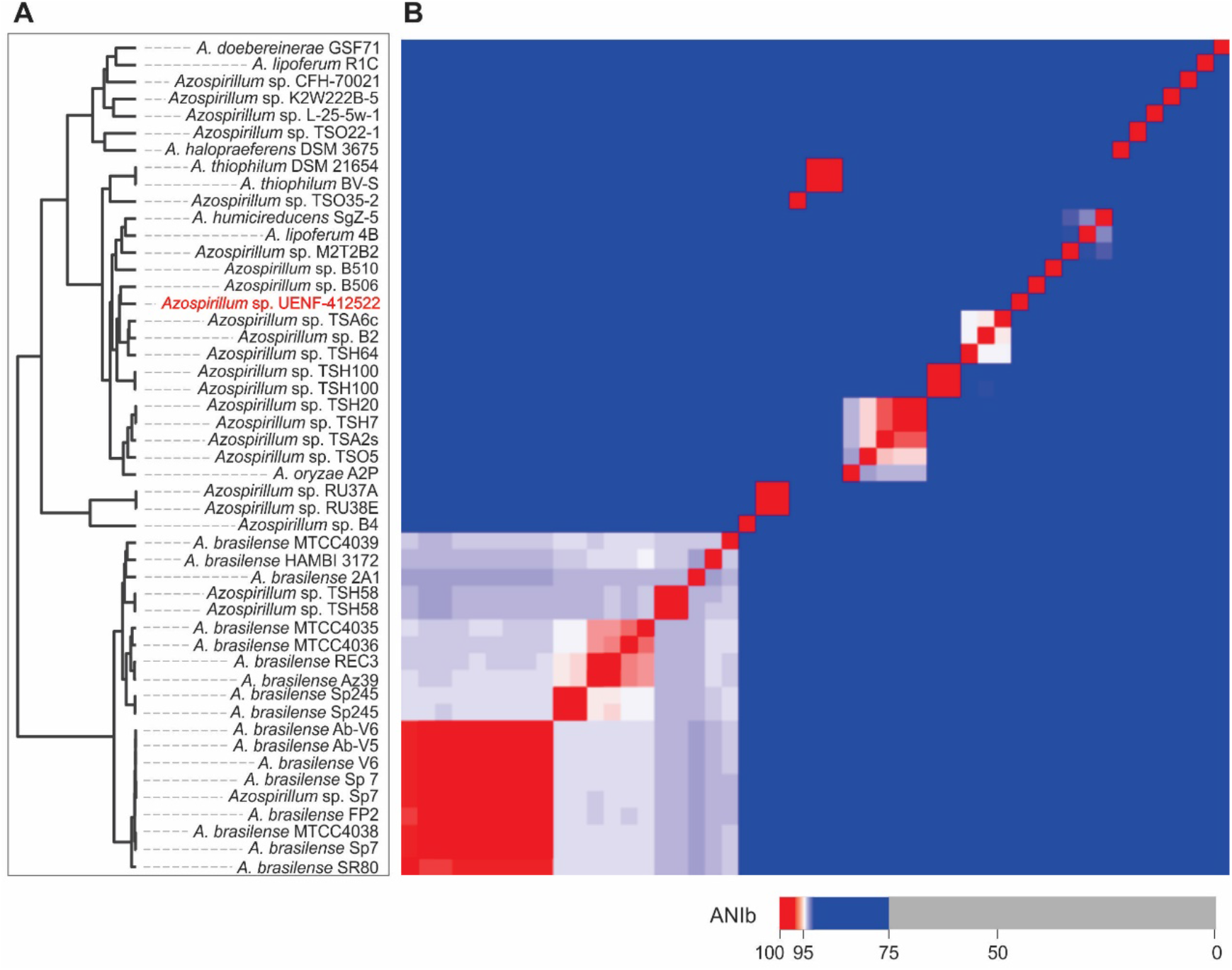
Average nucleotide identity (ANI). (A) Neighbor-joining cladogram. (B) Genome-to-genome distance matrix.

Even though the *Azospirillum* genus has more than 15 described species, genomic data are biased towards *A. brasilense*, which accounts for 37% (19 isolates) of the available genomes (Figure 3). We also noticed that five isolates have similar names and share more than 99.9% and 70% in ANI and dDDH analyses, respectively. We removed these redundant genomes from the downstream analyses and used GCF_001315015, GCF_003119195, GCF_003119115, and GCF_004923295 as representatives of *A. brasilense* Sp7, *A. brasilense* Sp245, *Azospirillum* sp. TSH58 and *Azospirillum* sp. TSH100 respectively. Hence, the dataset used in the next sections comprises 43 publicly available genomes and that of *Azospirillum* sp. UENF-412522 genome.

### Pangenome analyses of the *Azospirillum* genus

The pangenome is the complete gene repertoire of a given clade (e.g., a species) [28]. We employed a minimum identity threshold of 50% for gene family identification in *Azospirillum*. The pangenome comprises 42,515 genes, including a core genome (i.e. genes present in all isolates) of only 771 genes, likely because of the high diversity, prevalence of genomic rearrangements, and low synteny of this genus [9]. We clustered the isolates based on their gene presence/absence patterns and confirmed the high genomic heterogeneity across the genus (Figure 4A). Our results also support an open pangenome, as it grows with the addition of new genomes (Figure 4B). Similar results were previously shown in *Alphaproteobacteria* [29], who found 220 core genes across 27 in *Novosphingobium* spp. genomes. The *Azospirillum* core genome reported here is substantially greater than that, which could be at least partially explained by the fact that the vast majority of the isolates studied here were obtained from soil and plant tissues. *Azospirillum* sp. RU37 (GCF_900188305) and *Azospirillum* sp. RU38E (GCF_900188385) share the majority of accessory clusters. The same pattern is observed for *Azospirillum* sp. TSH20 (GCF_003115935) and *Azospirillum* sp. TSH7 (GCF_003115945). In both cases, the genomes with unusually high accessory genome similarity also exhibit 100% of ANI and dDDH, indicating that they belong to the same strain.

**Figure 4:**
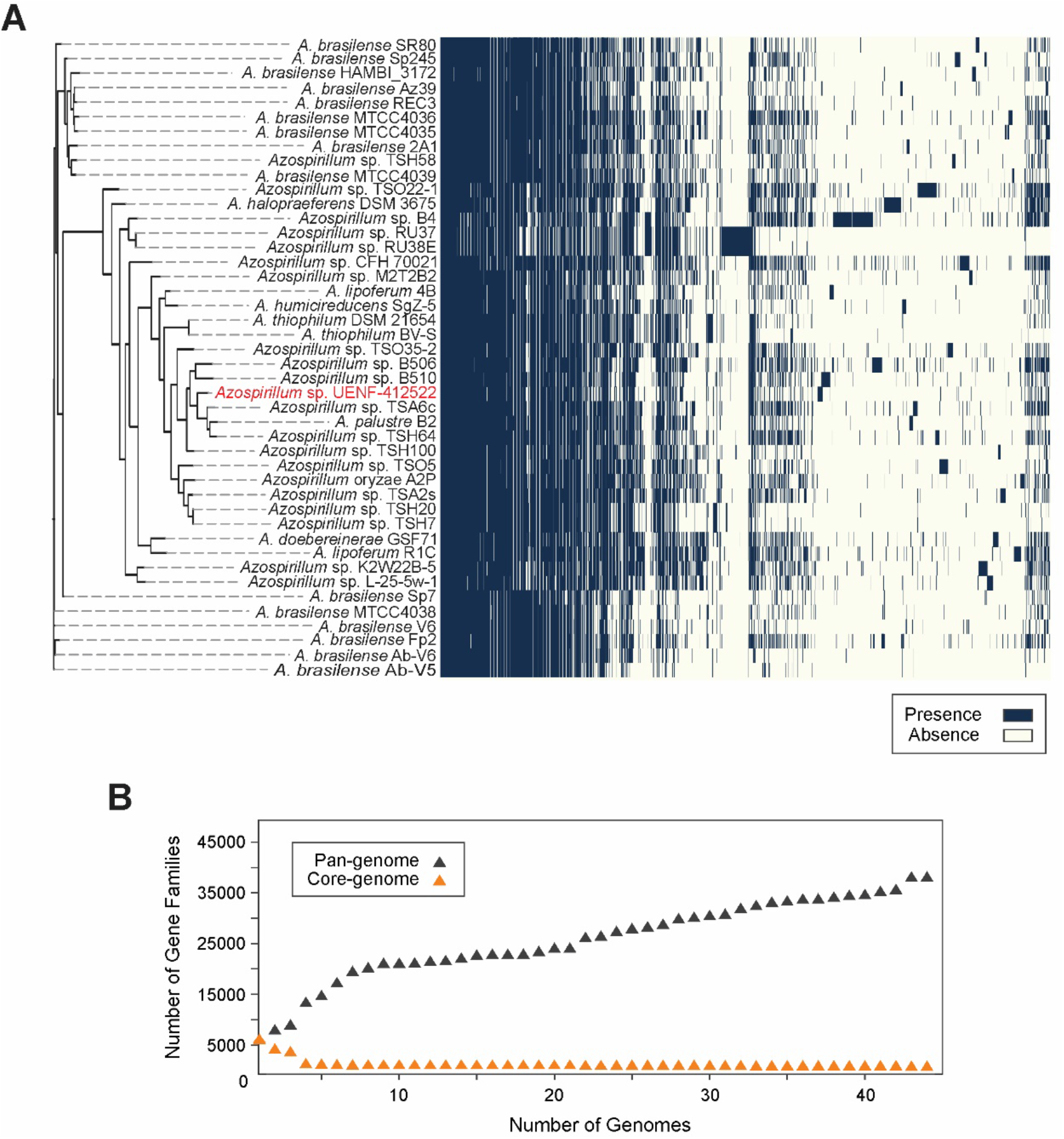
*Azospirillum* spp. pan-genome. (A) Gene presence/absence matrix, with gene presence depicted with solid blue boxes. (B) Pan- and core-genome sizes versus new genomes additions to the pan-genome dataset.

Next, we analyzed the unique gene complement of *Azospirillum* sp. UENF-412522. Out of the 18,065 unique genes inferred by Roary, 405 belonged to this strain. Among these genes, we identified creatinine amidohydrolase *crnA* (EI613_12750), an enzyme involved in the degradation of creatinine [30], which has been reported as replication-impairing molecule in bacteria [31]. The gene 3-alpha-hydroxysteroid dehydrogenase *hsdA* (EI613_09215), involved in steroid catabolism, is also exclusive to this strain and might assist *Azospirillum* sp. UENF-412522 to use steroids as a carbon source [32]. We also found the two component system *prsDE* (EI613_31600-31605) as exclusive to this strain. PrsDE produces a low molecular weight succinoglycan, which might assist in plant colonization by *Azospirillum sp.* UENF-412522 [33].

### Plant growth promotion genes

Plant growth modulation by bacteria is a complex process [34]. Several bacterial genes are often associated with direct beneficial effects, such as those responsible for biological nitrogen fixation (BFN), root growth enhancement, phosphate solubilization, and deviation of the ethylene biosynthesis pathway towards ammonia and α-ketobutyrate. We assessed the presence of genes that are likely involved in direct plant growth promotion across *Azospirillum* strains, with a particular focus in *Azospirillum sp.* UENF-412522. This procedure was guided by a maximum likelihood phylogenetic tree built with the protein sequences encoded by the core-genome, which was combined with a gene presence/absence matrix (Figure 5). Below we discuss these genes in light of the mechanisms by which they promote plant growth.

**Figure 5:**
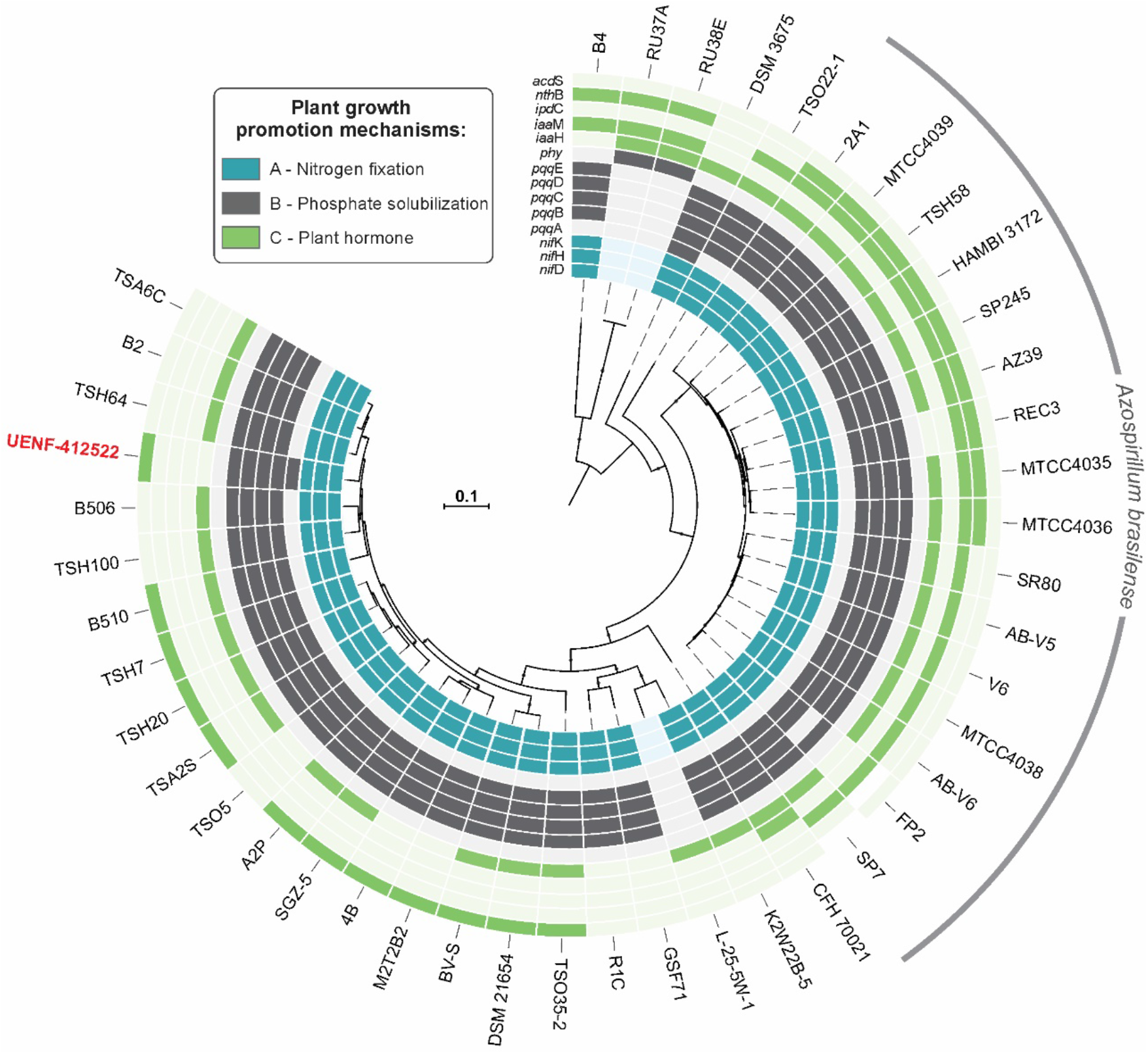
Core maximum likelihood phylogenetic tree. Plant growth-promoting genes were searched, grouped according to the mechanism of growth modulation, and mapped onto the tree.

BNF is a major plant growth promotion mechanism present in several *Azospirillum* strains [8, 35]. This process relies on the nitrogenase enzyme complex, which reduces atmospheric nitrogen (N_2_) to ammonium (NH_4_). The *nifHDK* genes encode, respectively, a nitrogenase iron (Fe) protein, a nitrogenase molybdenum-iron (MoFe) protein alpha chain, and a nitrogenase MoFe protein beta chain. The FeMo co-factor (FeMoCo) present in the MoFe protein binds N_2_, while the Fe protein uses the energy from ATP hydrolysis to drive the reduction of N_2_ to NH_4_ by FeMoCo [36]. We used the presence of the *nifHDK* genes to predict the nitrogen fixation capacity across *Azospirillum* genomes. All the analyzed genomes have these genes, except for *Azospirillum* sp. RU37A, *Azospirillum* sp. RU38E and *Azospirillum* sp. L-25-5w-1 (Figure 5). *Azospirillum* sp. UENF-412522 possesses the *nifHDK* (EI613_05225-05215) operon and a *nifDK* (EI613_05195-05200) operon, resembling the distribution found in most of the other *Azospirillum* genomes containing *nifHDK*. Some exceptions to this structure are found in *A. brasilense* Sp245, which possesses four *nifDK*, including vanadium nitrogenase variants [37]; *Azospirillum* sp. B506 presents a single *nifK*, and; *Azospirillum* sp. RU37A and *Azospirillum* sp. RU38E have two *nifD*. The presence of more than one *nifHDK* increase nitrogen fixation to the bacteria and associated plants. However, given that BNF is energetically expensive, the selective advantage of such increased activity remains unclear.

Most of the soil phosphate is insoluble and hence unavailable to direct plant absorption [38]. Several PGPRs promote plant growth by releasing organic acids that chelate divalent cations (e.g., Ca^2+^) in poorly soluble mineral forms (e.g., hydroxyapatite), increasing P availability [39]. Gluconic acid secretion is the best-characterized mechanism of phosphate solubilization, that is performed by pyrroloquinoline quinone (PQQ)-dependent glucose dehydrogenase (GDH) [40]. We used the presence of the genes encoding the PQQ co-factor as a proxy to understand P solubilization through GDH-PQQ in *Azospirillum* spp. The *pqq* genes found in *Azospirillum* sp. were *pqqABCDE* (EI613_19670-19690). While *Azospirillum* UENF-412522 and *A. halopraeferens* DSM3675 harbor the complete operon, all other strains lack *pqqA* (Figure 5). The *pqq* operon is absent in three strains (*Azospirillum* sp. RU37A, *Azospirillum* sp. RU38E, *Azospirillum* sp. L-25-5w-1). *A. brasilense* FP2 lacks *pqqA* and *pqqD*. Interestingly, phytase genes were only found in the pqq-lacking strains *Azospirillum* sp. RU37A and *Azospirillum* sp. RU38E, indicating that the capacity to mobilize P from organic compounds displaced PQQ-dependent P solubilization in these bacteria.

Although *pqqA* has been regarded as non-essential for PQQ biosynthesis [41], other studies support its role as the backbone for PQQ biogenesis [42]. PqqA binds to PqqD, prior to PqqAD interaction with PqqE [43]. The crucial role of *pqqA* and *pqqB* was also confirmed by knockout mutants of *Rahnella aquatilis* HX2, which showed decreased biocontrol and mineral P solubilization [44]. A direct correlation between *pqqB* and *pqqF* expression and PQQ production has also been shown [45]. Further, the same authors also reported that *Pseudomonas putida* KT2440 *pqqF* is controlled by an independent promoter and terminator, allowing this gene to modulate PQQ levels [45]. The presence of incomplete *pqq* operons in many *Azospirillum* strains allows one to speculate that these isolates have other genes to compensate for the absence of *pqqA*, a hypothesis that warrants experimental validation. Further, upstream of *pqqABCDE*, we found an alcohol dehydrogenase ADH IIB *qbdA* (EI613_19655) and its associated regulator *amgR* (EI613_19660), which are possibly related to acetic acid production mediated by PQQ [46]. This operon also has a a gene encoding a hypothetical protein containing a CXXCW motif, present in several soil bacteria, which might be associated with PQQ catabolism (TIGR03865). Hence, the *pqqABCDE* operon might be co-regulated with *qdbA* in *Azospirillum* sp. UENF-412522.

Ethylene is a plant hormone with a central role in senescence and growth of leaves, flowers, and fruits [47]. Under stress conditions, ethylene can impair plant cell elongation [48]. Ethylene is synthesized by the oxidation of 1-aminocyclopropane-1-carboxylate (ACC) by ACC oxidase. ACC can be broken down into α-ketobutyrate and NH_3_ by ACC deaminase [49], an enzyme produced by several PGPR that induce plant growth by lowering ethylene levels under stress conditions. We identified the ACC deaminase gene (*acdS*, EI613_03680) in *Azospirillum* sp. UENF-412522 and other 11 *Azospirillum* sp. strains, supporting that their association with plants can alleviate ethylene-mediated growth inhibition. Importantly, we have not found *acdS* in *A. brasilense* strains.

Considering its gene complement, *Azospirillum sp.* UENF-412522 has a significant potential to promote root growth. Despite the strong experimental evidence supporting IAA biosynthesis, we found none of the classic auxin biosynthesis genes (i.e., *ipdC*, *iaaH* and *iaaM*) [50–52], which prompted us to investigate other IAA biosynthesis pathways [51] in this strain. Interestingly, we found a gene encoding a nitrilase (*nit*, EI613_03300), which is 74% similar that encoded by a *nit* that converts indole-3-acetonitrile (IAN) directly into IAA or into indol-3-acetamide (IAM) in the rhizobacterium *Pseudomonas* sp. UW4 [53, 54]. In the latter pathway, the conversion of IAM into IAA requires the action of IAM-hydrolases, for which we found two candidate genes, EI613_04545 e EI613_07015. These genesare similar to amide hydrolases from *Rhodococcus* sp. (Q53116) [55, 56] and *Agrobacterium fabacearum* (ADY67766.1) [57], respectively. The apparent absence of tryptophan monooxygenase genes, which catalyze the conversion of Trp into IAM, indicates that *Azospirillum* sp. UENF 412522 produces IAA through an interplay between the IAN and IAM pathways, as reported in *Rhizobium* spp. and *Bradyrhizobium* spp. [53, 58]. Nevertheless, the initial steps in the production of IAN in *Azospirillum* sp. UENF 412522 are yet to be elucidated. In contrast, all *A. brasilense* strains possess the gene *ipdC* (Figure 5), supporting the ability of this species to synthesize IAA by the indole-3-pyruvate pathway, as experimentally demonstrated [59, 60]. Another eight *A. brasilense* strains possess nitrile hydratase (*nthB*) and *iaaH* genes, which contribute to IAA production by the conversion of IAN to IAM, and from IAM to IAA, respectively.

We also investigated *Azospirillum sp.* UENF-412522 genes that might promote root development through the emission of nitrite and nitric oxide. *A. brasilense* is known for secreting nitrite as a product of nitrate respiration [61, 62]. *Azospirillum sp.* UENF-412522 has a nitrate reductase operon *napCAB* (EI613_10540-10550). *napA* encodes a periplasmic nitrate reductase, while *napCB* encodes electron transfer subunits necessary for napA activity. This operon was proposed as an alternative electron acceptor for oxygen in *A. brasilense* sp. 245 periplasm [63] and is possibly responsible for nitrate respiration in *Azospirillum sp.* UENF-412522. The nitrite generated by nitrate respiration can be exported to the environment, absorbed by the plant root, and reduced to nitric oxide. Under acidic pH, nitrite can be reduced to nitric oxide without enzymatic action [64] and absorbed by root cells [65]. The downside of nitrate respiration to plant growth promotion is the reduction of available nitrogen in the system and the production of greenhouse gases. Hence, we hypothesize that *Azospirillum* sp. UENF-412522 nitrogen fixation contributes to nitrate respiration and plant absortion.

Most known *Azospirillum* strains are diazotrophs. N_2_-fixation requires a microaerophilic or anoxic environment because nitrogenase metalloclusters are sensitive to oxygen. Previous works in *Azospirillum* proposed that complete oxygen depletion inhibited nitrogenase activity [66], while others suggest otherwise [67]. *Azospirillum sp.* UENF-412522 was isolated from the rhizoplane, where low oxygen conditions are rather common [68]. Given the genomic investigations reported here, we hypothesize that P solubilization, root growth enhancement by nitrate respiration, and N_2_ fixation are dependent of an anoxic environment and might be interconnected in the soil-root system (Figure 6). Under low oxygen availability, nitrate is used as final electron acceptor in anaerobic respiration, producing nitrite, which is reduced to nitric oxide under low pH in the periplasm. Such acidic environment can be harnessed by the gluconic acid generated by the PQQ-dependent glucose dehydrogenase. Nitric oxide or nitrite would then be exported to the rhizosphere, stimulating root growth. Finally, gluconic acid is also exported to the rhizosphere, solubilizing phosphate for the root cells uptake.

**Figure 6:**
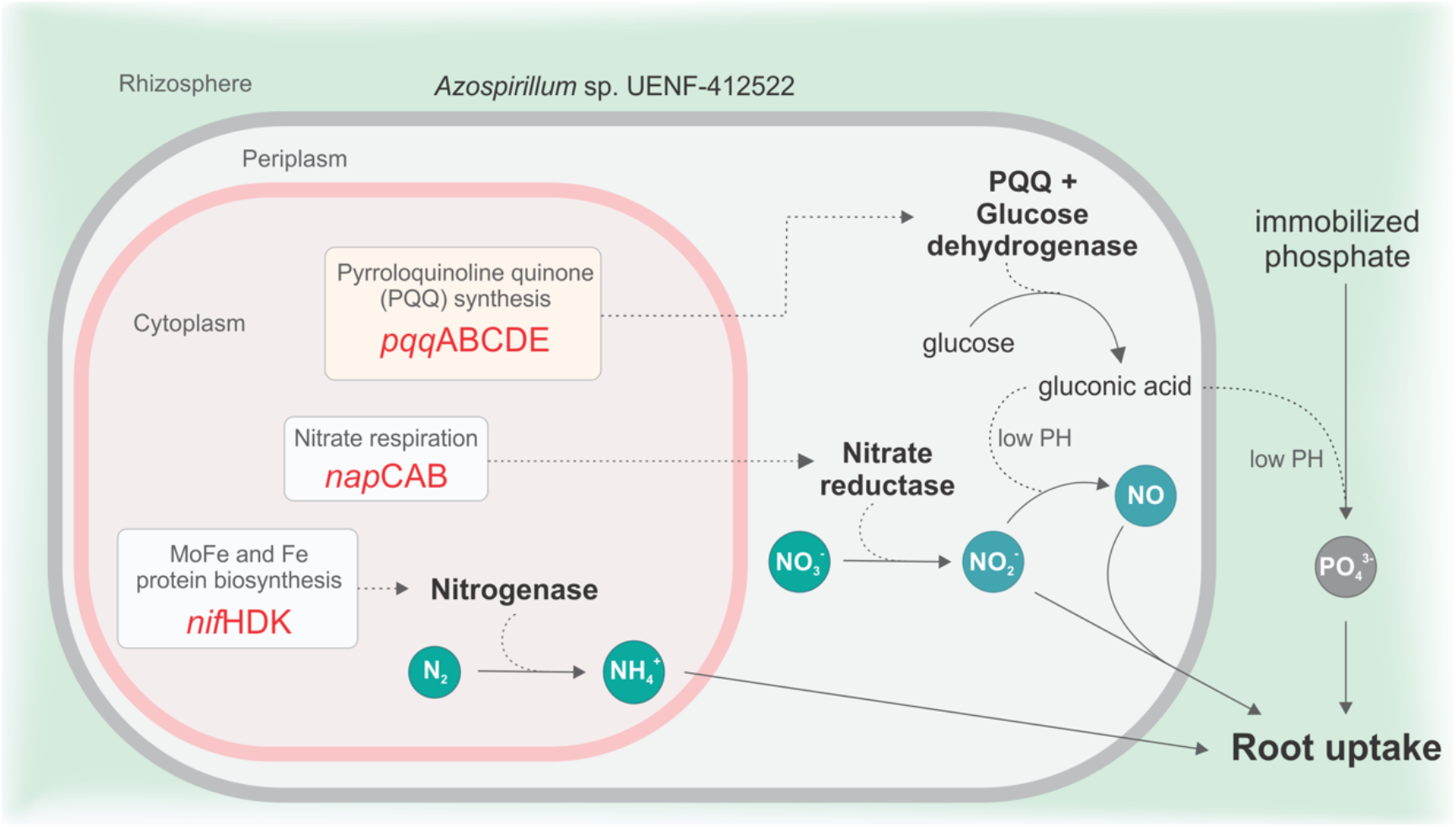
Schematic representation of nitrogen fixation, phosphate solubilization, and nitrite/nitric oxide emission by *Azospirillum sp.* UENF-412522 in the rhizosphere. The identified genes responsible for nitrogen fixation (*nifHDK*), root enhancement (*napCAB*), and phosphate solubilization (*pqqABCDE*) are represented within boxes. Nitrogen fixation occurs in the cytoplasm, while gluconic acid production and nitrate respiration take place at the periplasm. Ammonium generated by the nitrogenase can undergo anaerobic nitrification to nitrite, which can be exported to the periplasm. In the periplasm, glucose dehydrogenase, along with the PQQ co-factor, produces gluconic acid from glucose, lowering the pH. Nitrate is reduced to nitrite by a periplasmatic nitrate reductase napCAB. At low pH, nitrite can be reduced to nitric oxide. Acetic acid, nitrite, and nitric oxide can be exported to the rhizosphere. Acetic acid can reduce the pH in the rhizosphere and solubilize phosphate. The solubilized phosphate, nitrite, and nitric oxide can be captured by root cells.

### Antibiotic resistance genes

*Azospirillum sp.* UENF-412522 has several antibiotic resistance genes, such as those conferring resistance to: fosmidomycin (*fsr*, EI613_04505) [69], bicyclomycin (*bcr*, EI613_29470, EI613_14405) [70], bacitracin (*uppP/BacA*, EI613_27660) [71], tabtoxin (*ttr*, EI613_22745) [72], phenazine (*ehpR*, EI613_22490) [73], and tetracycline (*tetA*, EI613_26440) [74]. We also found multiple drug resistance genes (*mdtABCE*, EI613_19480-19470) [75], *acrAB* (EI613_25875-25880) [76], and *norM* (EI613_15640) [77]. Interestingly, most of these antibiotics, such as bicyclomycin [78], phenazine [79], bacitracin, and tetracycline [80], are commonly produced *Streptomyces*, which is abundant in the soil, where it plays a key role in plant organic matter decomposition [81, 82]. The presence of genes conferring resistance against *Streptomyces* antimicrobials is likely important to the survival of *Azospirillum sp.* UENF-412522 in the rhizosphere.

## CONCLUSION

In the present study we described *Azospirillum sp.* UENF-412522, a new *Azospirillum* plant growth-promoting species. We experimentally demonstrated that this bacterium is able to fix nitrogen, produce IAA, solubilize mineral P, and recognize and colonize seed-root surfaces. This strain is equipped with genes involved in BNF, P solubilization, ethylene catabolism, promotion of root development, and antibiotic resistance in the rhizosphere. Collectively, our results support the biotechnological potential of *Azospirillum sp.* UENF-412522 to be part of bioinoculant formulations for sustainable agriculture. We also propose a system involved in the partitioning of the fixed nitrogen between nitrate respiration and plant absorption, which is likely critical in plant-growth promotion under microaerophilic or anoxic conditions. Finally, this study emphasizes the importance of comprehensive studies using genomic, microscopy, microbiological, and biochemical approaches to characterize novel *Azospirillum* species outside of the *A. brasilense* clade, which could unlock the genus diversity and open novel possibilities for agricultural applications.

## METHODS

### Bacterial isolation and DNA purification

Passion fruit (*Passiflora edulis* f. *edulis*) root samples were obtained at the Integrated Agroecological Production System (SIPA), Embrapa Agrobiologia, Seropédica, Rio de Janeiro, Brazil (coordinates 22.7635°S 43.6886°W) and taken to the laboratory in a sterilized plastic bag kept in ice. Rhizoplane bacteria isolation was performed by an adapted version of Bramwell et al. (1995)[83]. In summary, root axis segments (10 g fresh weight) were sealed from both tip sides with paraffin, gently washed in tap water, and transferred to a 250 mL glass flask containing 90 mL of saline solution (NaCl, 0.9%). Roots were kept under agitation for 5 min at a rotatory shaker (200 rpm at 30 °C) to remove soil particles. After that, the same sealed root segment was carefully transferred to another 250 mL glass flask containing 90 mL of sterile saline (NaCl, 0.9%) solution plus 10 g of autoclaved sieved sand, where remained under agitation for 10 min at rotatory shaker (150 rpm at 30 °C) to detach bacteria cells from root surfaces (rhizoplane bacteria fraction). After sand decantation, 10 mL of the suspension were transferred to 90 mL of sterile saline solution (10^−1^ dilution), which were used to prepare successive dilutions until 10^−6^. Three independent 20 μL aliquots of all serial dilutions (10^−2^ to 10^−6^) were inoculated into glass vials (16 mL volume) containing 5 mL NFb nitrogen-free semisolid medium (composition in g L^−1^): malic acid, 5.0; K_2_HPO_4_, 0.5; MgSO_4_.7H_2_O, 0.2; NaCl, 0.1; CaCl_2_. 2H_2_O, 0.02; micronutrient stock solution (CuSO_4_.5H_2_O, 0.04; ZnSO_4_.7H_2_O, 0.12; H_3_BO_3_, 1.40; Na_2_MoO_4_.2H_2_O, 1.0; MnSO_4_. H_2_O, 1.175. Complete volume to 1,000 mL with distilled water), 2 mL; bromothymol blue (5 g L^−1^ in 0.2 N KOH), 2 mL; FeEDTA (solution 16.4 g L^−1^), 4 mL; vitamin stock solution (biotin, 10 mg; pyridoxal-HCl, 20 mg in 100 mL distilled water), 1 mL; KOH, 4.5 g; distilled water to bring the final volume to 1,000 mL and adjust pH to 6.5. A quantity of 1.6 g agar L^−1^ was added to prepare the semisolid medium [84].

Inoculated NFb vials were incubated in a growth chamber at 30 °C for 5 to 7 days. The growth of diazotrophic bacteria was characterized by the formation of a subsurface white pellicle. Using a loopful, a sample of the pellicle from the last positive dilution was inoculated in a fresh NFb N-free semisolid medium and incubated under the same conditions described above. This process was subsequently conducted for three times and the last new pellicle formed was streaked onto DYGS solid medium plates for colony purification. The DYGS composition (g L^−1^): glucose, 2.0; malic acid 2.0; peptone, 1.5; yeast extract, 2.0; K_2_HPO_4_, 0,5; MgSO_4_.7H_2_O, 0.5; glutamic acid, 1.5, pH is 6.5. The purified isolate was stored in 10% glycerol at −80 °C, under the code UENF-412522, as a diazotrophic bacterium associated with the passion fruit rhizoplane.

For DNA extraction, 100 μL of the stored bacteria were transferred to a glass tube containing 5 mL DYGS liquid medium and grown in an orbital shaker at 30 °C and 180 rpm for 48 hours. After that, 20 μL were transferred to the same medium and grown at the same conditions for 24 h. An aliquot of 200 μL was taken, and bacterial genomic DNA was extracted using the QIAamp kit (QIAGEN) following the manufacturer’s instructions. DNA was quantified in a 3000 NanoDrop (Thermo Scientific USA). DNA quality was analyzed using a 0.8% agarose gel stained with gel red.

To obtain an initial taxonomic assignment, PCR was conducted with the primers 27F (5‵-AGAGTTTGATCMTGGCTCAG 3‵) and 1492R (5’-TACGGYTACCTTGTTACGACTT-3‵). These reactions were performed using 50 ng of genomic DNA in 25 μL final reaction volume, containing: 2 μL of dNTPs (20 mM each) 2.5 μL of 10 x enzyme buffer, 0.75 μL 50 mM MgCl_2_, 2.5 μL of each primer (5 mM), 0.3 μL Taq polymerase (5 U/μL). Amplification was performed in a 96-well thermocycler Veriti model (Applied Biosystems), programmed at 9°C for 3 min, 30 cycles of amplification (94°C for 1 min, 55 °C for 30 s, 72 °C for 30 s) at 72 °C for 10 min. The sequencing reactions were performed using the Big Dye Terminator Sequencing Kit-Cycle Sequencing Ready ABI Prism version 3 (Life Technologies, USA), following manufacturer’s recommendations. Sequencing was performed on ABI Sequencer model 3130 (Applied Biosystems).

### Plant growth promotion traits of UENF-412522 bacterial strain

#### nifH gene detection

The *nifH* presence were assessed by PCR, with the primers PolF (5’TGCGAYCCSAARGCBGACTC 3‵) and PolR (5’ATSGCCATCATYTCRGCCGA 3’) [85]. In each reaction, we used 1 μL dNTPs (200 mM each), 5 μL of 10X buffer, 4 μL of 25 mM MgCl_2_, 0.5 μL of each primer (10 mM), 0.25 μL Taq polymerase (5 U/ μL) and 50 ng DNA, in a final volume of 50 μL. The amplification conditions were denaturation at 94 °C for 5 minutes, then 30 cycles (94 °C for 1 min, 55 °C for 1 min, 72 °C for 2 minutes) and a final extension step (72 °C for 5 min). PCR products were inspected on 1.5% agarose gels. Bp-100 DNA ladder (Invitrogen) was used as a molecular weight marker.

#### Acetylene reduction activity assay (ARA)

The *in vitro* nitrogen fixation ability of UENF-412522 was evaluated using the acetylene reduction assay (ARA), as previously described by Baldani [84]. The strain was grown in 16 mL glass vials containing 5 mL of N-free NFB semisolid medium, as described above. These flasks were inoculated with 20 μL of the bacterial inoculum suspended in sterile water, adjusted to an optical density (OD) of 1.0, and incubated at 30 °C for 48 h. After pellicle formation, the vials were closed with a sterilized pierceable rubber stopper of the subseal type. Syringes were used to remove 1 mL of air and to inject 1 mL of acetylene in each vial. The flasks were incubated at 30 °C for 1 h, and 1 mL of the gas phase was analyzed on a gas chromatograph with flame ionization (Perkin Elmer), to determine the ethylene concentration in the sample.

#### Phosphorus and zinc solubilization

Phosphate solubilization was independently carried out using 1 g.L^−1^ Araxá rock phosphate [Ca_10_(PO_4_)_6_F_2_] and tricalcium phosphate [Ca_3_(PO_4_)_2_]. Bacterial cultures grown in DYGS liquid medium adjusted to an optical density of 1.0 were inoculated (20 μL) at the center of the petri dish with agar containing Pikovskaya medium. After this procedure, they were incubated at 30 °C for 7 days. The ability of the isolates in solubilizing P was evaluated by measuring the translucent halo, according to Kumar and Narula formula [86], in which S.I. (solubility index) = halo Diameter (mm)/colony diameter (mm). We conducted three biological replicates of this assay.

We also assessed P solubilization in Pikovskaya liquid medium supplemented with 1g.L^−1^ of the Araxá rock phosphate or Ca_10_(PO_4_)_6_F_2_ was used to quantify the soluble P. Aliquots of 100 μL of bacterial cultures grown in DYGS liquid medium were transferred to 50 mL tubes containing Pikovskaya medium and kept for 7 days under constant agitation at 150 rpm in a rotatory shaker at 30 °C. After this period, cultures were centrifuged at 3200 rpm for 15 minutes, and the supernatant was used for pH determination and quantification of soluble P.

Zinc solubilization was carried out according to Intorne et al [87]. Aliquots of 20 μL of bacterial strains cultivated for 24 h at 30 °C in DYGS liquid medium, adjusted to an optical density of 1.0, were inoculated at the center of the petri dish containing Saravanan medium with agar supplemented with 1g.L^−1^ of ZnO and incubated at 30 °C for 7 days. After this period, the ability to solubilize ZnO was also performed according to Kumar and Narula [86]. We conducted three biological replicates of this assay.

#### Indole acetic acid (IAA) production

Bacterial isolates were cultured in liquid DYGS medium for 24 h at 30 °C and transferred (25 μL) to test tubes containing 5 mL of the same medium in the presence and absence of L-tryptophan (100 mg.L^−1^) before incubation in the dark for 72 h at 30 °C, shaking at 150 rpm. The cultures were then transferred to 2 mL tubes and centrifuged at 10,000 rpm for 10 min, and the supernatant transferred to a test tube with 2 mL of Salkowski reagent [88]. Tubes were incubated for 30 min in the dark. The production of IAA was evaluated by the presence of pink color in the tubes, and the color intensity was determined with a spectrophotometer at a wavelength of 530 nm. IAA concentrations were measured using a calibration curve.

#### Biochemical API 50 CH/E test

For the metabolic characterization of bacterial isolates, we used the API 50 CH/E test kit (bioMérieux SA Marcy-I’Etoile/France), which evaluates the ability of the isolates to conduct fermentation of 49 carbohydrates and derivatives. To this end, the isolates were cultured in DYGS medium for 24 h at 150 rpm. After this period, the isolates were suspended in autoclaved water and adjusted to optical density 1.0; 2 mL of this suspension were added to the CHL medium (bioMérieux) and transferred to galleries containing different substrates. Each dome was sealed with a sterile drop of mineral oil and incubated in an environmental chamber at 37 °C. Reads were recorded after 24 h and 48 h after inoculation, and eventual production of organic acids during incubation shifts the pH indicator from red to yellow.

### Plant growth promotion assay under greenhouse conditions

A greenhouse assay was carried out to evaluate the effect of UENF-412522 inoculation in plants. To perform it, passion fruit seeds of the cultivar Yellow master FB200 were disinfected in 1% sodium hypochlorite solution for 20 minutes, followed by three rinses in autoclaved distilled water and then transferred to germitest paper moistened with autoclaved water. Then, the seeds were incubated in a growth chamber at 30 °C, for 16 h in the light and 8 h in the dark. After germinating, these seeds were transferred to styrofoam trays containing autoclaved plant substrate under greenhouse conditions. After 30 days of transplanting, seedlings of approximately 6 cm were transferred to a 300 mL plastic bag containing the same autoclaved plant growth substrate. The inoculum of the UENF-412522 strain was prepared in liquid DYGS medium after 40 h of growth in a rotatory shaker at 30 °C and 150 rpm. Bacterial suspensions containing approximately 10^8^ cells.mL^−1^ (O.D. = 1.0 at 600 ηm) were inoculated on the region of the plant’s neck, and plants were transferred to the greenhouse for 30 days. The assay was set up in a randomized block design, two treatments (inoculated and non-inoculated plantlets) and six replications. The control plants received 1 mL of the liquid DYGS medium containing lysed UENF-412522 cells.

### Plant root colonization assay

We evaluated plant-bacteria interaction by determining the population size and root colonization by scanning electron microscopy (SEM). For this, passion fruit seeds of the Yellow cultivar master FB200 were disinfected and placed to germinate as described above. After germination, the seedlings were transferred to a test tube containing autoclaved vermiculite. Then, 1 ml of bacterial suspension containing approximately 10^8^ cells mL^−1^ was inoculated on the seedling neck. Seedlings were kept for seven days after inoculation (d.a.i) in a culture room under a temperature of 30°C (16 h in the light and 8 hours in the dark). For SEM, root samples were collected at 1 and 7 days after the inoculation of the bacterial suspension. Samples for SEM were prepared according to Baldotto et al. [89] and viewed at Zeiss DSEM 962 at 15 Kv voltage in secondary electron detector mode. The bacteria count at rhizosphere, rhizoplane, and roots was performed at seven d.a.i., as described above. The control consisted of seedlings inoculated with the DYGS medium.

### Genome sequencing and assembly

The sequencing procedures and basic data processing were performed, as previously described [25]. In summary, paired-end libraries were prepared using the TruSeq Nano DNA LT Library Prep (Illumina) and sequenced on a HiSeq 2500 instrument at the Life Sciences Core Facility (LaCTAD; UNICAMP, Campinas, Brazil). Sequencing reads (2 × 100 bp) had their quality checked with FastQC 0.11.5 (https://www.bioinformatics.babraham.ac.uk/projects/fastqc/). Quality filtering was performed with Trimmomatic 0.35 [90], and reads with average quality below 30 were discarded. The genome assembly was performed using SPAdes 3.8 [91] and assembly metrics evaluated by QUAST 3.0 [92].

### Genome annotation and phylogenetic analysis

The assembled genome was annotated with the NCBI Prokaryotic Genome Annotation Pipeline [93]. The *Azospirillum* sp. UENF-412522 genome was deposited on Genbank under the BioProject PRJNA508400. *Azospirillum* spp. genomes available in RefSeq [94, 95] were downloaded (n = 48, as of June 03 2019) and protein sequences were predicted using PROKKA 1.13.3 [96]. We searched for proteins encoded by genes potentially involved in plant growth promotion using a blastp [97] search on the SwissProt database, with minimum coverage and similarity thresholds of 70% and 60%, respectively. All-against-all average nucleotide identity (ANI) was computed using pyani 0.28 [98]. Pan-genome analysis was performed with Roary 3.12.0, with a minimum identity threshold of 50% [99]. Multiple sequence alignments were performed using Muscle 3.8.30 [100] and phylogenetic reconstructions performed with RAxML 8.2.10 [101].

## AUTHOR CONTRIBUTIONS

Conceived the study: FLO, TMV; Funding and resources: FLO, TMV; Data analysis: GLR, FPM, RKG, FP-S, DC-A, IP-O; Performed experiments: PSLG, STS, AFA; Interpretation of the results: GLR, FPM, FLO, TMV; Wrote the manuscript: GLR, FPM, FLO, TMV.

## ACKNOWLEDGEMENTS

This work was supported by Fundação Carlos Chagas Filho de Amparo à Pesquisa do Estado do Rio de Janeiro (FAPERJ; grants E-26/111.827/2013, E-26/102.259/2013, E-26/201.239/2014, E-26/203.309/2016 and E-26/203.014/2018), Coordenação de Aperfeiçoamento de Pessoal de Nível Superior - Brasil (CAPES; Finance Code 001), and Conselho Nacional de Desenvolvimento Científico e Tecnológico (CNPq; grant 449904/2014-8 and 314263/2018-7). The funding agencies had no role in the design of the study and collection, analysis, and interpretation of data and in writing.

## REFERENCES

1. Bourguet D, Guillemaud T: The hidden and external costs of pesticide use. In: Sustainable Agriculture Reviews. Springer; 2016: 35–120.

2. Maroniche GA, Diaz PR, Borrajo MP, Valverde CF, Creus CM: Friends or foes in the rhizosphere: traits of fluorescent Pseudomonas that hinder Azospirillum brasilense growth and root colonization. FEMS Microbiol Ecol 2018, 94(12).

3. Vejan P, Abdullah R, Khadiran T, Ismail S, Nasrulhaq Boyce A: Role of Plant Growth Promoting Rhizobacteria in Agricultural Sustainability-A Review. Molecules 2016, 21(5).

4. Prasad M, Srinivasan R, Chaudhary M, Choudhary M, Jat LK: Plant Growth Promoting Rhizobacteria (PGPR) for Sustainable Agriculture: Perspectives and Challenges. In: PGPR Amelioration in Sustainable Agriculture. Elsevier; 2019: 129–157.

5. Brink SC: Unlocking the Secrets of the Rhizosphere. Trends Plant Sci 2016, 21(3):169–170.

6. Ahkami AH, White III RA, Handakumbura PP, Jansson C: Rhizosphere engineering: Enhancing sustainable plant ecosystem productivity. Rhizosphere 2017, 3:233–243.

7. Mehnaz S: Azospirillum: a biofertilizer for every crop. In: Plant Microbes Symbiosis: Applied Facets. Springer; 2015: 297–314.

8. Fukami J, Cerezini P, Hungria M: Azospirillum: benefits that go far beyond biological nitrogen fixation. AMB Express 2018, 8(1):73.

9. Wisniewski-Dye F, Borziak K, Khalsa-Moyers G, Alexandre G, Sukharnikov LO, Wuichet K, Hurst GB, McDonald WH, Robertson JS, Barbe V et al: Azospirillum genomes reveal transition of bacteria from aquatic to terrestrial environments. PLoS Genet 2011, 7(12):e1002430.

10. Tarrand JJ, Krieg NR, Dobereiner J: A taxonomic study of the Spirillum lipoferum group, with descriptions of a new genus, Azospirillum gen. nov. and two species, Azospirillum lipoferum (Beijerinck) comb. nov. and Azospirillum brasilense sp. nov. Can J Microbiol 1978, 24(8):967–980.

11. Reis VM, Baldani VLD, Baldani JI: Isolation, Identification and Biochemical Characterization of Azospirillum spp. and Other Nitrogen-Fixing Bacteria. In: Handbook for Azospirillum: Technical Issues and Protocols. Edited by Cassán FD, Okon Y, Creus CM. Cham: Springer International Publishing; 2015: 3–26.

12. Pankievicz VC, do Amaral FP, Santos KF, Agtuca B, Xu Y, Schueller MJ, Arisi AC, Steffens MB, de Souza EM, Pedrosa FO et al: Robust biological nitrogen fixation in a model grass-bacterial association. Plant J 2015, 81(6):907–919.

13. Rodriguez H, Gonzalez T, Goire I, Bashan Y: Gluconic acid production and phosphate solubilization by the plant growth-promoting bacterium Azospirillum spp. Naturwissenschaften 2004, 91(11):552–555.

14. Cohen AC, Bottini R, Pontin M, Berli FJ, Moreno D, Boccanlandro H, Travaglia CN, Piccoli PN: Azospirillum brasilense ameliorates the response of Arabidopsis thaliana to drought mainly via enhancement of ABA levels. Physiol Plant 2015, 153(1):79–90.

15. Garcia JE, Maroniche G, Creus C, Suarez-Rodriguez R, Ramirez-Trujillo JA, Groppa MD: In vitro PGPR properties and osmotic tolerance of different Azospirillum native strains and their effects on growth of maize under drought stress. Microbiol Res 2017, 202:21–29.

16. D’Angioli AM, Viani RAG, Lambers H, Sawaya ACHF, Oliveira RS: Inoculation with Azospirillum brasilense (Ab-V4, Ab-V5) increases Zea mays root carboxylate-exudation rates, dependent on soil phosphorus supply. Plant and soil 2017, 410(1-2):499–507.

17. Vacheron J, Renoud S, Muller D, Babalola OO, Prigent-Combaret C: Alleviation of abiotic and biotic stresses in plants by Azospirillum. In: Handbook for Azospirillum. Springer; 2015: 333–365.

18. Cassán F, Diaz-Zorita M: Azospirillum sp. in current agriculture: From the laboratory to the field. Soil Biology and Biochemistry 2016, 103:117–130.

19. Hungria M, Ribeiro RA, Nogueira MA: Draft Genome Sequences of Azospirillum brasilense Strains Ab-V5 and Ab-V6, Commercially Used in Inoculants for Grasses and Legumes in Brazil. Genome Announc 2018, 6(20).

20. Blaha D, Prigent-Combaret C, Mirza MS, Moënne-Loccoz Y: Phylogeny of the 1-aminocyclopropane-1-carboxylic acid deaminase-encoding gene acdS in phytobeneficial and pathogenic Proteobacteria and relation with strain biogeography. FEMS Microbiology Ecology 2006, 56(3):455–470.

21. Esquivel-Cote R, Ramírez-Gama RM, Tsuzuki-Reyes G, Orozco-Segovia A, Huante PJP, Soil: Azospirillum lipoferum strain AZm5 containing 1-aminocyclopropane-1-carboxylic acid deaminase improves early growth of tomato seedlings under nitrogen deficiency. 2010, 337(1):65–75.

22. Vikram A, Alagawadi AR, Krishnaraj PU, Mahesh Kumar KSJWJoM, Biotechnology: Transconjugation studies in Azospirillum sp. negative to mineral phosphate solubilization. 2007, 23(9):1333–1337.

23. Kaneko T, Minamisawa K, Isawa T, Nakatsukasa H, Mitsui H, Kawaharada Y, Nakamura Y, Watanabe A, Kawashima K, Ono A et al: Complete genomic structure of the cultivated rice endophyte Azospirillum sp. B510. DNA Res 2010, 17(1):37–50.

24. Malhotra M, Srivastava S: An ipdC gene knock-out of Azospirillum brasilense strain SM and its implications on indole-3-acetic acid biosynthesis and plant growth promotion. Antonie Van Leeuwenhoek 2008, 93(4):425–433.

25. Matteoli FP, Passarelli-Araujo H, Reis RJA, da Rocha LO, de Souza EM, Aravind L, Olivares FL, Venancio TM: Genome sequencing and assessment of plant growth-promoting properties of a Serratia marcescens strain isolated from vermicompost. BMC Genomics 2018, 19(1):750.

26. Matteoli FP, Passarelli-Araujo H, Pedrosa-Silva F, Olivares FL, Venancio TM: Population structure and pangenome analysis of Enterobacter bugandensis uncover the presence of blaCTX-M-55, blaNDM-5 and blaIMI-1, along with sophisticated iron acquisition strategies. Genomics 2020, 112(2):1182–1191.

27. Eckert B, Weber OB, Kirchhof G, Halbritter A, Stoffels M, Hartmann A: Azospirillum doebereinerae sp. nov., a nitrogen-fixing bacterium associated with the C4-grass Miscanthus. Int J Syst Evol Microbiol 2001, 51(Pt 1):17–26.

28. Xiao J, Zhang Z, Wu J, Yu J: A brief review of software tools for pangenomics. Genomics Proteomics Bioinformatics 2015, 13(1):73–76.

29. Kumar R, Verma H, Haider S, Bajaj A, Sood U, Ponnusamy K, Nagar S, Shakarad MN, Negi RK, Singh Y et al: Comparative Genomic Analysis Reveals Habitat-Specific Genes and Regulatory Hubs within the Genus Novosphingobium. mSystems 2017, 2(3).

30. Beuth B, Niefind K, Schomburg D: Crystal structure of creatininase from Pseudomonas putida: a novel fold and a case of convergent evolution. J Mol Biol 2003, 332(1):287–301.

31. McDonald T, Drescher KM, Weber A, Tracy S: Creatinine inhibits bacterial replication. J Antibiot (Tokyo) 2012, 65(3):153–156.

32. Mobus E, Maser E: Molecular cloning, overexpression, and characterization of steroid-inducible 3alpha-hydroxysteroid dehydrogenase/carbonyl reductase from Comamonas testosteroni. A novel member of the short-chain dehydrogenase/reductase superfamily. J Biol Chem 1998, 273(47):30888–30896.

33. York GM, Walker GC: The Rhizobium meliloti exoK gene and prsD/prsE/exsH genes are components of independent degradative pathways which contribute to production of low-molecular-weight succinoglycan. Mol Microbiol 1997, 25(1):117–134.

34. Numan M, Bashir S, Khan Y, Mumtaz R, Shinwari ZK, Khan AL, Khan A, Al-Harrasi A: Plant growth promoting bacteria as an alternative strategy for salt tolerance in plants: A review. Microbiol Res 2018, 209:21–32.

35. Steenhoudt O, Vanderleyden J: Azospirillum, a free-living nitrogen-fixing bacterium closely associated with grasses: genetic, biochemical and ecological aspects. FEMS Microbiol Rev 2000, 24(4):487–506.

36. Dixon R, Kahn D: Genetic regulation of biological nitrogen fixation. Nat Rev Microbiol 2004, 2(8):621–631.

37. Wisniewski-Dye F, Lozano L, Acosta-Cruz E, Borland S, Drogue B, Prigent-Combaret C, Rouy Z, Barbe V, Herrera AM, Gonzalez V et al: Genome Sequence of Azospirillum brasilense CBG497 and Comparative Analyses of Azospirillum Core and Accessory Genomes provide Insight into Niche Adaptation. Genes (Basel) 2012, 3(4):576–602.

38. Ewel JJ, Schreeg LA, Sinclair TR: Resources for Crop Production: Accessing the Unavailable. Trends Plant Sci 2019, 24(2):121–129.

39. Sharma SB, Sayyed RZ, Trivedi MH, Gobi TA: Phosphate solubilizing microbes: sustainable approach for managing phosphorus deficiency in agricultural soils. Springerplus 2013, 2:587.

40. Alori ET, Glick BR, Babalola OO: Microbial Phosphorus Solubilization and Its Potential for Use in Sustainable Agriculture. Front Microbiol 2017, 8:971.

41. Toyama H, Lidstrom ME: pqqA is not required for biosynthesis of pyrroloquinoline quinone in Methylobacterium extorquens AM1. Microbiology 1998, 144 (Pt 1):183–191.

42. Barr I, Latham JA, Iavarone AT, Chantarojsiri T, Hwang JD, Klinman JP: Demonstration That the Radical S-Adenosylmethionine (SAM) Enzyme PqqE Catalyzes de Novo Carbon-Carbon Cross-linking within a Peptide Substrate PqqA in the Presence of the Peptide Chaperone PqqD. J Biol Chem 2016, 291(17):8877–8884.

43. Latham JA, Iavarone AT, Barr I, Juthani PV, Klinman JP: PqqD is a novel peptide chaperone that forms a ternary complex with the radical S-adenosylmethionine protein PqqE in the pyrroloquinoline quinone biosynthetic pathway. J Biol Chem 2015, 290(20):12908–12918.

44. Li L, Jiao Z, Hale L, Wu W, Guo Y: Disruption of gene pqqA or pqqB reduces plant growth promotion activity and biocontrol of crown gall disease by Rahnella aquatilis HX2. PLoS One 2014, 9(12):e115010.

45. An R, Moe LA: Regulation of Pyrroloquinoline Quinone-dependent glucose dehydrogenase activity in the model rhizosphere-dwelling bacterium Pseudomonas putida KT2440. Appl Environ Microbiol 2016, 82(16):4955–4964.

46. Yakushi T, Matsushita K: Alcohol dehydrogenase of acetic acid bacteria: structure, mode of action, and applications in biotechnology. Appl Microbiol Biotechnol 2010, 86(5):1257–1265.

47. Iqbal N, Khan NA, Ferrante A, Trivellini A, Francini A, Khan MIR: Ethylene Role in Plant Growth, Development and Senescence: Interaction with Other Phytohormones. Front Plant Sci 2017, 8:475.

48. Vaseva, II, Qudeimat E, Potuschak T, Du Y, Genschik P, Vandenbussche F, Van Der Straeten D: The plant hormone ethylene restricts Arabidopsis growth via the epidermis. Proc Natl Acad Sci U S A 2018, 115(17):E4130–E4139.

49. Glick BR: Bacteria with ACC deaminase can promote plant growth and help to feed the world. Microbiol Res 2014, 169(1):30–39.

50. Spaepen S, Vanderleyden J, Remans R: Indole-3-acetic acid in microbial and microorganism-plant signaling. FEMS Microbiol Rev 2007, 31(4):425–448.

51. Spaepen S, Vanderleyden J: Auxin and plant-microbe interactions. Cold Spring Harb Perspect Biol 2011, 3(4).

52. Imada EL, Rolla Dos Santos AA, Oliveira AL, Hungria M, Rodrigues EP: Indole-3-acetic acid production via the indole-3-pyruvate pathway by plant growth promoter Rhizobium tropici CIAT 899 is strongly inhibited by ammonium. Res Microbiol 2017, 168(3):283–292.

53. Duca D, Rose DR, Glick BR: Characterization of a nitrilase and a nitrile hydratase from Pseudomonas sp. strain UW4 that converts indole-3-acetonitrile to indole-3-acetic acid. Appl Environ Microbiol 2014, 80(15):4640–4649.

54. Duca DR, Rose DR, Glick BR: Indole acetic acid overproduction transformants of the rhizobacterium Pseudomonas sp. UW4. Antonie Van Leeuwenhoek 2018, 111(9):1645–1660.

55. Mayaux JF, Cerbelaud E, Soubrier F, Yeh P, Blanche F, Petre D: Purification, cloning, and primary structure of a new enantiomer-selective amidase from a Rhodococcus strain: structural evidence for a conserved genetic coupling with nitrile hydratase. J Bacteriol 1991, 173(21):6694–6704.

56. Patten CL, Blakney AJ, Coulson TJ: Activity, distribution and function of indole-3-acetic acid biosynthetic pathways in bacteria. Crit Rev Microbiol 2013, 39(4):395–415.

57. Delamuta JRM, Scherer AJ, Ribeiro RA, Hungria M: Genetic diversity of Agrobacterium species isolated from nodules of common bean and soybean in Brazil, Mexico, Ecuador and Mozambique, and description of the new species Agrobacterium fabacearum sp. nov. Int J Syst Evol Microbiol 2020, 70(7):4233–4244.

58. Vega-Hernández MC, León-Barrios M, Pérez-Galdona R: Indole-3-acetic acid production from indole-3-acetonitrile in Bradyrhizobium. Soil Biology and Biochemistry 2002, 34(5):665–668.

59. Vande Broek A, Lambrecht M, Eggermont K, Vanderleyden J: Auxins upregulate expression of the indole-3-pyruvate decarboxylase gene in Azospirillum brasilense. J Bacteriol 1999, 181(4):1338–1342.

60. El-Khawas H, Adachi KJB, Soils Fo: Identification and quantification of auxins in culture media of Azospirillum and Klebsiella and their effect on rice roots. 1999, 28(4):377–381.

61. Creus CM, Graziano M, Casanovas EM, Pereyra MA, Simontacchi M, Puntarulo S, Barassi CA, Lamattina L: Nitric oxide is involved in the Azospirillum brasilense-induced lateral root formation in tomato. Planta 2005, 221(2):297–303.

62. Molina-Favero C, Creus CM, Simontacchi M, Puntarulo S, Lamattina L: Aerobic nitric oxide production by Azospirillum brasilense Sp245 and its influence on root architecture in tomato. Mol Plant Microbe Interact 2008, 21(7):1001–1009.

63. Steenhoudt O, Keijers V, Okon Y, Vanderleyden J: Identification and characterization of a periplasmic nitrate reductase in Azospirillum brasilense Sp245. Arch Microbiol 2001, 175(5):344–352.

64. Bethke PC, Badger MR, Jones RL: Apoplastic synthesis of nitric oxide by plant tissues. Plant Cell 2004, 16(2):332–341.

65. Molina-Favero C, Mónica Creus C, Lanteri ML, Correa-Aragunde N, Lombardo MC, Barassi CA, Lamattina L: Nitric Oxide and Plant Growth Promoting Rhizobacteria: Common Features Influencing Root Growth and Development. In: Advances in Botanical Research. vol. 46: Academic Press; 2007: 1–33.

66. Zhang Y, Burris RH, Ludden PW, Roberts GP: Regulation of nitrogen fixation in Azospirillum brasilense. FEMS Microbiol Lett 1997, 152(2):195–204.

67. Malinich EA, Bauer CE: Transcriptome analysis of Azospirillum brasilense vegetative and cyst states reveals large-scale alterations in metabolic and replicative gene expression. Microb Genom 2018, 4(8).

68. Lecomte SM, Achouak W, Abrouk D, Heulin T, Nesme X, Haichar FeZ: Diversifying Anaerobic Respiration Strategies to Compete in the Rhizosphere. 2018, 6(139).

69. Fujisaki S, Ohnuma S, Horiuchi T, Takahashi I, Tsukui S, Nishimura Y, Nishino T, Kitabatake M, Inokuchi H: Cloning of a gene from Escherichia coli that confers resistance to fosmidomycin as a consequence of amplification. Gene 1996, 175(1-2):83–87.

70. Bentley J, Hyatt LS, Ainley K, Parish JH, Herbert RB, White GR: Cloning and sequence analysis of an Escherichia coli gene conferring bicyclomycin resistance. Gene 1993, 127(1):117–120.

71. El Ghachi M, Bouhss A, Blanot D, Mengin-Lecreulx D: The bacA gene of Escherichia coli encodes an undecaprenyl pyrophosphate phosphatase activity. J Biol Chem 2004, 279(29):30106–30113.

72. Anzai H, Yoneyama K, Yamaguchi I: The nucleotide sequence of tabtoxin resistance gene (ttr) of Pseudomonas syringae pv. tabaci. Nucleic Acids Res 1990, 18(7):1890.

73. Giddens SR, Feng Y, Mahanty HK: Characterization of a novel phenazine antibiotic gene cluster in Erwinia herbicola Eh1087. Mol Microbiol 2002, 45(3):769–783.

74. Waters SH, Rogowsky P, Grinsted J, Altenbuchner J, Schmitt R: The tetracycline resistance determinants of RP1 and Tn1721: nucleotide sequence analysis. Nucleic Acids Res 1983, 11(17):6089–6105.

75. Nagakubo S, Nishino K, Hirata T, Yamaguchi A: The putative response regulator BaeR stimulates multidrug resistance of Escherichia coli via a novel multidrug exporter system, MdtABC. J Bacteriol 2002, 184(15):4161–4167.

76. Okusu H, Ma D, Nikaido H: AcrAB efflux pump plays a major role in the antibiotic resistance phenotype of Escherichia coli multiple-antibiotic-resistance (Mar) mutants. J Bacteriol 1996, 178(1):306–308.

77. Brown MH, Paulsen IT, Skurray RA: The multidrug efflux protein NorM is a prototype of a new family of transporters. Mol Microbiol 1999, 31(1):394–395.

78. Vior NM, Lacret R, Chandra G, Dorai-Raj S, Trick M, Truman AW: Discovery and Biosynthesis of the Antibiotic Bicyclomycin in Distantly Related Bacterial Classes. Appl Environ Microbiol 2018, 84(9).

79. Abdelfattah MS, Ishikawa N, Karmakar UK, Yamaku K, Ishibashi M: New phenazine analogues from Streptomyces sp. IFM 11694 with TRAIL resistance-overcoming activities. J Antibiot (Tokyo) 2016, 69(6):446–450.

80. Procopio RE, Silva IR, Martins MK, Azevedo JL, Araujo JM: Antibiotics produced by Streptomyces. Braz J Infect Dis 2012, 16(5):466–471.

81. Chater KF, Biro S, Lee KJ, Palmer T, Schrempf H: The complex extracellular biology of Streptomyces. FEMS Microbiol Rev 2010, 34(2):171–198.

82. Chater KF: Recent advances in understanding Streptomyces. F1000Res 2016, 5:2795.

83. Bramwell PA, Barallon RV, Rogers HJ, Bailey MJ: Extraction and PCR amplification of DNA from the rhizoplane. In: Molecular microbial ecology manual. Springer; 1995: 89–108.

84. Baldani JI, Reis VM, Videira SS, Boddey LH, Baldani VLD: The art of isolating nitrogen-fixing bacteria from non-leguminous plants using N-free semi-solid media: a practical guide for microbiologists. Plant and Soil 2014, 384(1):413–431.

85. Poly F, Ranjard L, Nazaret S, Gourbiere F, Monrozier LJ: Comparison of nifH gene pools in soils and soil microenvironments with contrasting properties. Appl Environ Microbiol 2001, 67(5):2255–2262.

86. Kumar V, Narula N: Solubilization of inorganic phosphates and growth emergence of wheat as affected by Azotobacter chroococcum mutants. Biology and Fertility of Soils 1999, 28(3):301–305.

87. Intorne AC, de Oliveira MV, Lima ML, da Silva JF, Olivares FL, de Souza Filho GA: Identification and characterization of Gluconacetobacter diazotrophicus mutants defective in the solubilization of phosphorus and zinc. Arch Microbiol 2009, 191(5):477–483.

88. Sarwar M, Kremer RJ: Determination of bacterially derived auxins using a microplate method. Letters in Applied Microbiology 1995, 20(5):282–285.

89. Baldotto LEB, Olivares FL, Bressan-Smith R: Structural interaction between GFP-labeled diazotrophic endophytic bacterium Herbaspirillum seropedicae RAM10 and pineapple plantlets ‘Vitória’. Brazilian Journal of Microbiology 2011, 42:114–125.

90. Bolger AM, Lohse M, Usadel B: Trimmomatic: a flexible trimmer for Illumina sequence data. Bioinformatics 2014, 30(15):2114–2120.

91. Bankevich A, Nurk S, Antipov D, Gurevich AA, Dvorkin M, Kulikov AS, Lesin VM, Nikolenko SI, Pham S, Prjibelski AD et al: SPAdes: a new genome assembly algorithm and its applications to single-cell sequencing. J Comput Biol 2012, 19(5):455–477.

92. Gurevich A, Saveliev V, Vyahhi N, Tesler G: QUAST: quality assessment tool for genome assemblies. Bioinformatics 2013, 29(8):1072–1075.

93. Tatusova T, DiCuccio M, Badretdin A, Chetvernin V, Nawrocki EP, Zaslavsky L, Lomsadze A, Pruitt KD, Borodovsky M, Ostell J: NCBI prokaryotic genome annotation pipeline. Nucleic acids research 2016, 44(14):6614–6624.

94. O’Leary NA, Wright MW, Brister JR, Ciufo S, Haddad D, McVeigh R, Rajput B, Robbertse B, Smith-White B, Ako-Adjei D et al: Reference sequence (RefSeq) database at NCBI: current status, taxonomic expansion, and functional annotation. Nucleic Acids Res 2016, 44(D1):D733–745.

95. Pruitt KD, Tatusova T, Maglott DR: NCBI reference sequences (RefSeq): a curated non-redundant sequence database of genomes, transcripts and proteins. Nucleic Acids Res 2007, 35(Database issue):D61–65.

96. Seemann T: Prokka: rapid prokaryotic genome annotation. Bioinformatics 2014, 30(14):2068–2069.

97. Altschul SF, Madden TL, Schaffer AA, Zhang J, Zhang Z, Miller W, Lipman DJ: Gapped BLAST and PSI-BLAST: a new generation of protein database search programs. Nucleic Acids Res 1997, 25(17):3389–3402.

98. Pritchard L, Glover RH, Humphris S, Elphinstone JG, Toth IK: Genomics and taxonomy in diagnostics for food security: soft-rotting enterobacterial plant pathogens. Analytical Methods 2016, 8(1):12–24.

99. Page AJ, Cummins CA, Hunt M, Wong VK, Reuter S, Holden MT, Fookes M, Falush D, Keane JA, Parkhill J: Roary: rapid large-scale prokaryote pan genome analysis. Bioinformatics 2015, 31(22):3691–3693.

100. Edgar RC: MUSCLE: multiple sequence alignment with high accuracy and high throughput. Nucleic Acids Res 2004, 32(5):1792–1797.

101. Stamatakis A: RAxML version 8: a tool for phylogenetic analysis and post-analysis of large phylogenies. Bioinformatics 2014, 30(9):1312–1313.

