## Supplementary figures and table S1 for "Characterization of cellular, biochemical and genomic features of the diazotrophic plant growth-promoting bacterium *Azospirillum* sp. UENF-412522, a novel member of the *Azospirillum* genus"

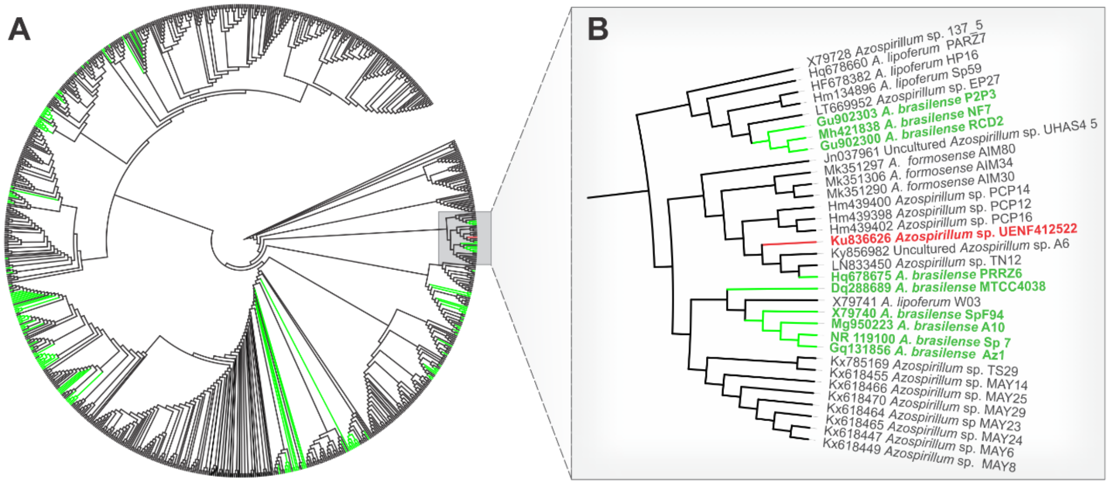


**Figure S1:** 16S rRNA maximum likelihood phylogenetic reconstruction of *Azospirillum* spp. sequences. (A) Tree overview of all 16S rRNA sequences available in Genbank (n = 894 as of June 2019). Green branches represent *A. brasilense* sequences. (B) Close view of the clade containing *Azospirillum* sp. UENF-412522 (in red), which also comprise isolates of three other species (i.e. *A. lipoferum*, *A. brasilense,* and *A. formosense*).

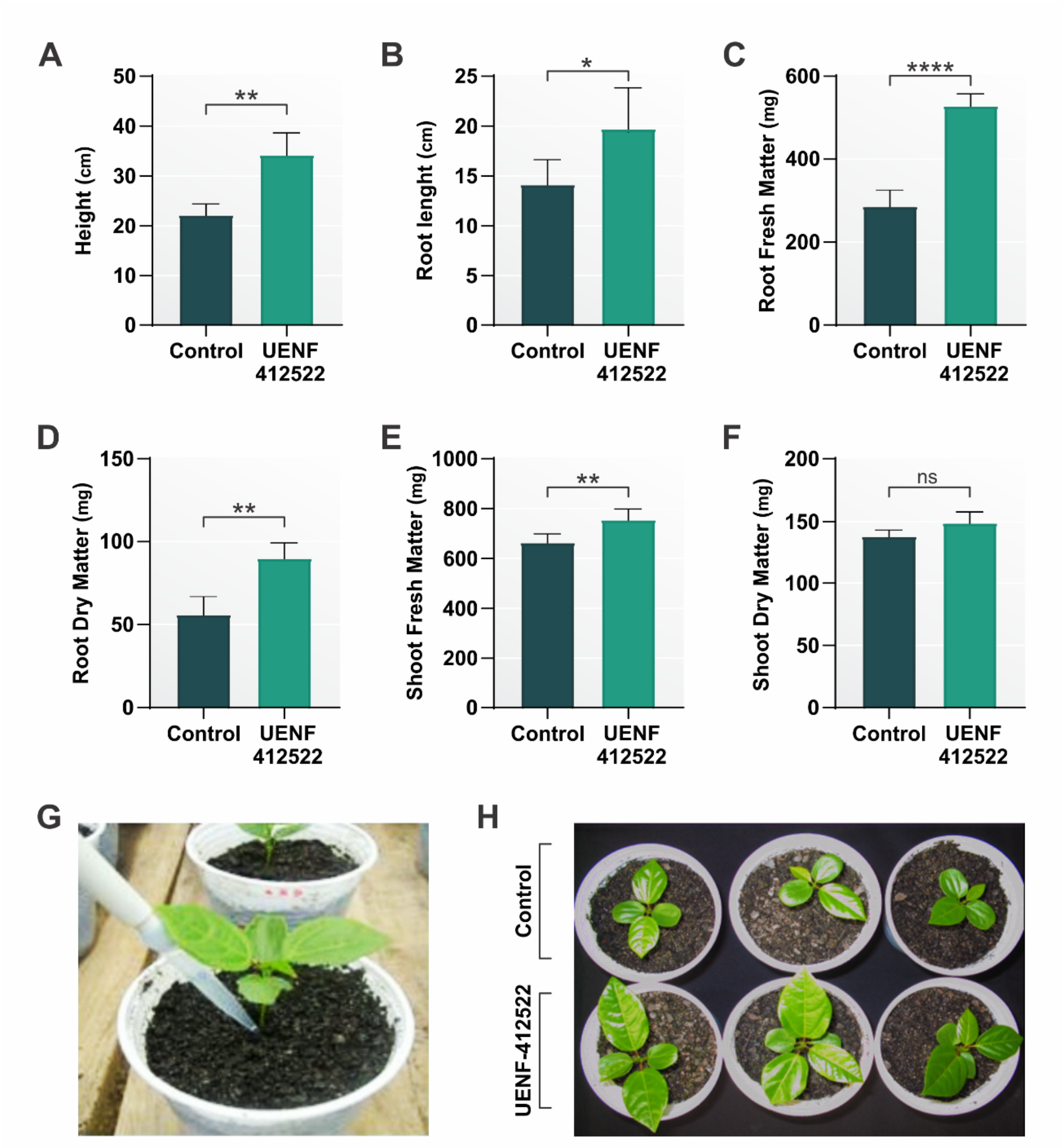


**Figure S2:** In vivo effects of *Azospirillum* sp. UENF-412622 inoculation on passion fruit plantlets growth. (A) Shoot height. (B) Root length. (C) Root fresh matter. (D) Root dry matter. (E) Shoot fresh matter. (G) Inoculation with the bacteria suspension. (H) Typical biostimulation effect observed after inoculation with *Azospirillum* sp. UENF-412622. Statistics (ANOVA): *: $p$-value <0.05; **: $p$-value <0.01; ***: $p$-value <0.001; ****: $p$-value <0.0001.

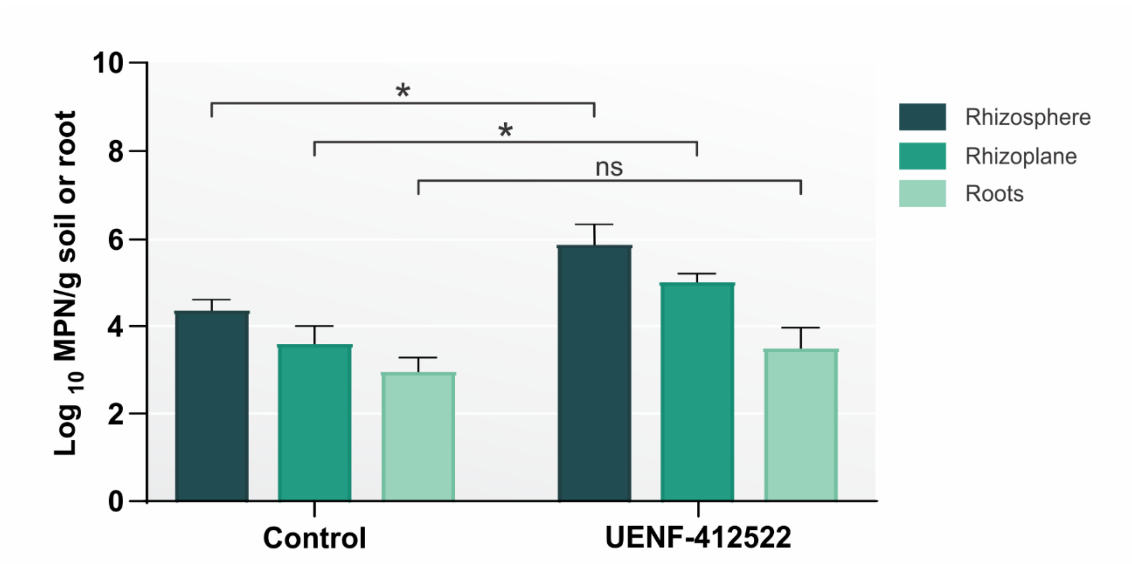


**Figure S3:** Estimation of diazotrophic bacteria population associated to rhizosphere soil, rhizoplane and roots, under gnotobiotic condition of inoculated passion fruit plantlets. Control plants showed natural diazotrophic bacteria population. Data expressed in Most Probable Number (MPN) of cell transformed in $\log_{10}$ per g of soil (rhizosphere) or root (rhizoplane or root). Statistics (ANOVA): *: *p*-value <0.05; **: *p*-value <0.01; ***: *p*-value <0.001; ****: *p*-value <0.0001.

**Table S1:** API 50 CH system carbon utilizations profile for *Azospirillum sp. UENF-412522, Azospirillum lipoferum* ATCC 29707T/DSM 1691T, *Azospirillum doebereinerae* DSM 13131T and *Azospirillum brasilense* ATTC 29145T/ DSM1690T. The + and – symbols stand for ability and inability to metabolize a carbon source, respectively.

| | ***Azospirillum* sp. UENF-412522** | ***Azospirillum lipoferum* ATCC 29707[1]** | ***Azospirillum doebereinerae* DSM 13131[2]** | ***Azospirillum brasilense* ATTC 29145[3]** |
|---|---|---|---|---|
| **Non-distinctive C-sources** | | | | |
| Glycerol | + | + | + | + |
| L-arabinose | + | + | + | + |
| D-galactose | + | + | + | + |
| D-glucose | + | + | + | + |
| D-fructose | + | + | + | + |
| L-rhamnose | - | - | - | - |
| Inositol | - | - | - | - |
| D-maltose | - | - | - | - |
| D-saccharose | - | - | - | - |
| **Distinctive C-sources** | | | | |
| D-ribose | + | + | - | - |
| D-mannitol | - | + | + | - |
| D-sorbitol | - | + | + | - |
| N-acetylglucosamine | - | + | - | + |

[1]*Azospirillum lipoferum* ATCC 29707T/DSM 1691T (strain 59b; Tarrand et al.) [2]*Azospirillum doebereinerae* (Eckert et al.) strain type DSM 13131T (strain GSF71T) [3]*Azospirillum brasilense* ATTC 29145T/DSM1690T (strain Sp7), isolated from grass rhizosphere soil in Rio de Janeiro

### REFERENCES

Eckert B, Weber OB, Kirchhof G, Halbritter A, Stoffels M, Hartmann A: Azospirillum doebereinerae sp. nov., a nitrogen-fixing bacterium associated with the C4-grass Miscanthus. Int J Syst Evol Microbiol 2001, 51(Pt 1):17-26.

Tarrand JJ, Krieg NR, Dobereiner J: A taxonomic study of the Spirillum lipoferum group, with descriptions of a new genus, Azospirillum gen. nov. and two species, Azospirillum lipoferum (Beijerinck) comb. nov. and Azospirillum brasilense sp. nov. Can J Microbiol 1978, 24(8):967-980.
